# A Novel Approach Towards Less Invasive Multi ‘Omics Gut Analyses: A Pilot Study

**DOI:** 10.1101/2021.11.24.469968

**Authors:** Adam J. Berlinberg, Ana Brar, Andrew Stahly, Mark E. Gerich, Blair P. Fennimore, Frank I. Scott, Kristine A. Kuhn

**Affiliations:** Division of Rheumatology, Department of Medicine, University of Colorado – Anschutz Medical Campus, Aurora, CO; Department of Medicine, University of Colorado – Anschutz Medical Campus, Aurora, CO; Division of Gastroenterology and Hepatology, Department of Medicine, University of Colorado – Anschutz Medical Campus, Aurora, CO

## Abstract

Newer ‘omics approaches such as metatranscriptomics and metabolomics allow functional assessments of the interaction(s) between the gut microbiome and the human host. In order to generate meaningful data with these approaches, though, the method of sample collection is critical. Prior studies have relied upon expensive and invasive means towards sample acquisition such as intestinal biopsy, while other studies have relied upon easier methods of collection such as fecal samples that do not necessarily represent those microbes in contact with the host. In this pilot study, we attempt to characterize a novel, minimally invasive method towards sampling the human microbiome using mucosal cytology brush sampling compared to intestinal gut biopsy on 5 healthy participants undergoing routine screening colonoscopy. We compared metatranscriptomic analyses between the two collection methods, identifying increased taxonomic evenness and beta diversity in the cytology brush samples, and similar community transcriptional profiles between the two methods. Metabolomics assessment demonstrated striking differences between the two methods, implying a difference in bacterial-derived versus human absorbed metabolites. Put together, this study supports the use of a less invasive method of microbiome sampling with cytology brushes, but caution must be exercised when performing metabolomics assessment as this represents differential metabolite production but not absorption by the host.

**Importance:** In order to generate meaningful metabolomic and microbiome data, the method of sample collection is critical. This study utilizes and compares two methods to intestinal tissue collection for evaluation of metabolites and microbiome, finding that using a brush to sample the microbiome is superior to tissue biopsy. However, for metabolomics assessment, biopsy may still be required.

## Introduction

The human intestinal microbiome is colonized by trillions of commensal microorganisms (1), and alterations in microbiome composition, or dysbiosis, associate with a wide variety of inflammatory, metabolic, and infectious human diseases(2-4). The combination of metagenomic, metatranscriptomic, and metabolomic data together with 16S rRNA taxonomic profiling enhances microbial community data, providing functional insights into the role of the human gut microbiome in disease(5). Essential to such multi ‘omics techniques are representative sample collection that can be obtained simply, efficiently, and at low cost, yet provide high-quality data.

The geographic landscape of microbial colonization in the intestine varies based upon micro-anatomic site (6, 7). Thus, studies of the gut microbiome require consideration with regards to the collection method as a difference between mucosal versus luminal inhabitants exists, and different collection methods consequently affect results. A number of various methods have been utilized in the literature such as fecal sampling, tissue biopsy, or endoscopic brushing, each with advantages and disadvantages as highlighted in a recent review article(8). The simplest and routinely performed collection method involves fecal sampling, although some argue that it does not represent the true mucosal microbiome(9, 10) sampled from the outer mucosal layer where most microbes inhabit (11, 12). In addition to luminal bacteria that do not have contact with the host as well as sloughed bacteria from the mucus layer, fecal samples contain a significant amount of host genetic material, thus complicating some types of analyses like metagenomics and metatranscriptomics. Less frequently will the more gold standard technique of pinch biopsy be used due to cost, invasiveness, and subject discomfort. Prior studies have utilized pinch biopsies and have identified greater microbial diversity with significant differences in diversity analyses compared to fecal samples(13).

Despite the available collection methods, the fundamental challenge remains finding a method of sampling that is non-invasive, cost-effective, and provides an accurate depiction of the mucosal microbial environment. To address these issues, we propose a method using a cytology brush inserted in the rectum to brush the luminal surface. The collection can be performed in the clinic and requires minimal equipment. No sedation or bowel preparation is required, and the procedure can be performed in a few minutes, thus minimizing discomfort for patients. Furthermore, the cost savings are significant compared to endoscopy and aspiration capsule. Described here is a pilot study demonstrating the feasibility of this method compared to endoscopic biopsy controls. We find the cytology brush method increased evenness and beta diversity compared to endoscopic biopsy sampling, though metabolomic analysis demonstrates a differential metabolite profile indicative of the diverse mucosal layer including bacterial and human derived metabolites that differs from the deeper biopsy samples.

## Methods

### Sample Collection

Five healthy individuals were recruited through the University of Colorado gastroenterology (GI) clinic between December 2019 and February 2020. Participants were identified from the endoscopy schedule as undergoing routine screening colonoscopy. Inclusion criteria included any subject over the age of 18 undergoing routine colonoscopy. Exclusion criteria included use of antiplatelet drugs or chronic anticoagulation, use of immunomodulatory medications, non-steroidal anti-inflammatory drugs seven days before colonoscopy, active malignancy, decompensated cirrhosis, chronic kidney disease on dialysis, or history of inflammatory bowel disease. Patients were identified from the colonoscopy schedule, and written informed consent was obtained just prior to the procedure. This study was conducted according to the principles within the Declaration of Helsinki. All study procedures were approved by the Colorado Multiple Institutional Review Board (protocol #14-2012).

Patients prepared for colonoscopy by standard protocol the night prior. Following conscious sedation per routine care, one cytology brush (Fisher #22-281660) was inserted 3 cm beyond the anal verge, pressed against the lateral wall, and rotated two full turns. The brush was then placed into 500 μL RNAlater (ThermoFisher) in a 1.5 mL Eppendorf tube and stored on ice. A second cytology brush was then inserted 3 cm beyond the anal verge against the opposite lateral wall, rotated two times, then placed in 500 μL PBS on ice. Lastly, routine colonoscopy was performed with four pinch biopsies obtained at approximately the same depth as the cytology brushes. One biopsy sample was placed in RNAlater and the remaining three samples in PBS and then placed on ice. All samples were then taken to the laboratory and frozen at -80°C immediately.

### Microbial RNA Isolation, Library Prep, and Metatranscriptomics Sequencing

RNA was isolated using the AllPrep Power Fecal DNA/RNA kit (Qiagen). For initial input, brushes in RNAlater were vortexed at maximum speed for 15 seconds and 200 μL of this suspension was used. The manufacturer’s protocol was then followed with the exception of incubating samples for 15 minutes in the presence of lysis buffer and 25 μL DTT prior to removal of solid tissue for the biopsy samples, followed by homogenization of samples for both methods in the same manner. Quality control was performed using a Thermo Scientific NanoDrop 2000 spectrophotometer ensuring 260/280 nm light ratios >1.8 for all samples. Libraries were then constructed utilizing 5-10 ng RNA for each sample and using the Next Ultra II Directional RNA Library Prep Kit with rRNA depletion (New England Bioscience) in a paired end fashion with 2×150 base pair paired end reads. Libraries underwent quality control via tape station prior to multiplexing at a concentration of 4 nM, and sequencing was performed on an Illumina MiSeq platform (San Diego, CA, USA) at the University of Colorado Genomics core with >6 Gb data output per sample.

### Data Processing and Taxonomic Analysis

Manual inspection of sequenced reads was performed utilizing FastQC v0.11.9 for all samples. Paired end reads were then concatenated and quality control conducted with Kneaddata 0.7.5 (http://huttenhower.sph.harvard.edu/kneaddata), utilizing Trimmomatic v0.39(14) and Bowtie2 v2.3.5(15) to remove unwanted human genome reads and low quality sequences. The processed reads were then entered into the HUMAnN 2.0 pipeline(16), utilizing MetaPhlAn v2.0(17) which does not account for paired-end relationships, with gene profiling abundance performed using the UniRef90 full universal database. Output data in reads per kilobase was then converted to copies per million prior to downstream application utilizing the command: humann2_renorm_table. Alpha diversity was determined on a species level using MicrobiomeAnalyst (18, 19) with the Observed, Chao, Shannon, and Simpson methods. Beta diversity between the two groups was assessed utilizing Bray-Curtis dissimilarity and visualized with a Principal Coordinate Analysis (PCoA) plot. PERMANOVA was performed with MicrobiomeAnalyst of the beta diversity clustering.

### Functional Analysis

HUMAnN 2.0 was used with default settings to obtain gene family abundance for each sample individually prior to combining and normalizing based upon sequencing depth. Analysis was performed after renaming normalized gene families to Kyoto Encyclopedia of Genes and Genomes (KO) pathways (humann2_regroup_table). Metatranscriptomic abundance was assessed using the functional diversity profile on MicrobiomeAnalyst and top pathways identified through read abundance.

### Metabolomics

Metabolomics analysis by LC-MS of tissue collected by cytology brush versus biopsies was performed by the University of Colorado Metabolomics Core. Cytology brush samples in PBS were spun at 4°C for 10 min at 18 213 rcf. Then the brush was removed and centrifugation repeated. PBS was aspirated and replaced with 100 uL of ice cold 5:3:2 MeOH:MeCN:water (v/v/v). For biopsy tissue, samples were centrifuged twice for 10 minutes at 18 213 rcf at 4°C, and then PBS aspirated and replaced with 700 µL of MeOH:MeCN:water. Extracted samples were vortexed for 30 min at 4 °C, centrifuged once as before, and then an aliquot of supernatant was transferred to an autosampler vial for analysis. Samples were analyzed on a Thermo Vanquish UHPLC coupled to a Thermo Q Exactive mass spectrometer. Metabolites were separated on a 5 min C18 gradient with positive and negative (separate runs) electrospray ionization. Data acquisition and analysis was performed as previously described(20, 21). Quality control was assessed using technical replicates injected every 10 runs. Resulting .raw files were converted to .mzXML format using RawConverter then metabolites assigned and peak areas integrated using Maven (Princeton University) in conjunction with the KO database and an in-house standard library of >600 compounds. The targeted data analysis focused on metabolites involved in central carbon and nitrogen metabolism and yielded measurements of 114 metabolites. No post hoc normalization was performed; data is available upon request. Samples were normalized relative to each other based upon the same initial starting weight of tissue.

### Data Analysis

Taxonomic and metatranscriptomic profiling was performed using MicrobiomeAnalyst software. For taxonomy alpha diversity, Student’s t-test was utilized using the methods of Chao, Shannon, and Simpson as well as the observed species index as they passed Shapiro-Wilk normality tests. Beta diversity was assessed utilizing Bray-Curtis dissimilarity and visualized with a Principal Coordinate Analysis (PCoA) plot. Relative abundance OTU differences were compared using Wilcoxon signed-rank tests and adjusted with an FDR 1%. PERMANOVA was performed using MicrobiomeAnalyst software of the beta diversity clustering. Metabolomics assessment was performed using MetaboAnalyst software following log transformation(22). Statistical assessments were done with an ANOVA with Kruskal-Wallis post-hoc test or paired t-test where noted.

## Results

### Cytology Brush Sampling Provides Improved Bacterial DNA Recovery and Microbial Diversity

The primary goal of this study was to evaluate a novel less-invasive method for microbiome sampling utilizing a cytology brush in contrast to the gold standard colon pinch biopsy. Therefore, we compared the bacterial microbiome in paired cytology brushes to tissue biopsy in five individuals undergoing standard-of-care cancer screening colonoscopies. In all, an average of 20.7 million paired end reads was performed per sample. Paired end reads were concatenated and Kneaddata was utilized to remove low quality and human-genome derived reads. After quality control, a total of 14.5±7.2 million reads remained in the biopsy samples and 41.7±31.9 million reads in the paired cytology brush samples, of which 7.7±5.1 million reads remained after removal of human reads in the biopsy group, and 36.3±29.8 million reads in the cytology brush group (Supplemental Tables 1 and 2). Of these, a total of 26.5±11.5% of filtered reads correlated with bacterial sequences in the biopsies, compared to 59.1±23.9% of reads in the brushes (Figure 1A; p=0.0256 by unpaired t-test). After removal of human reads, the vast majority of remaining sequences aligned to bacterial reads in both sample types that was used for downstream analysis (96.1±2.9% in the biopsy samples, and 97.9±3.5% in the cytology brush samples), with the remaining reads of viral etiology discarded. As predicted, these data support the brush as a superior collection method for bacterial DNA compared to colon biopsies.

**Figure 1.**
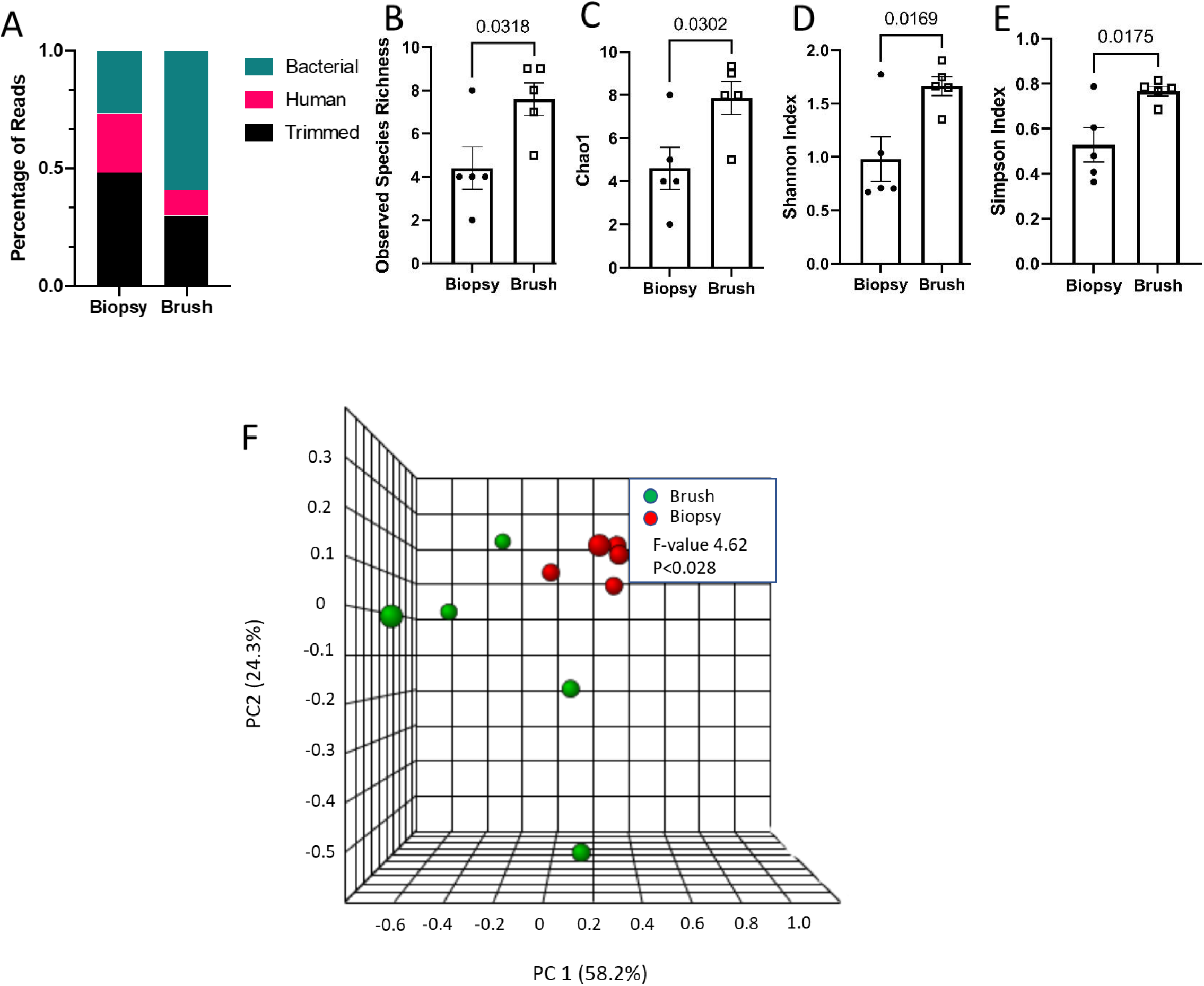
Cytology brush sampling, compared to biopsies, results in higher abundance of bacteria-derived sequence reads and increased diversity. Metatrascriptomic analysis of bacterial communities was performed on paired cytology brush sampling and colon pinch biopsies from five healthy participants undergoing colonoscopy. (A) Percentage of sequencing reads broken down in terms of bacterial, human, and trimmed reads between sample types. (B-E) Alpha diversity calculated in MicrobiomeAnalyst using the methods of (B) observed species richness, (C) Chao1, (D) Shannon, and (E) Simpson. Values for each subject are shown as a symbol and bars represent the group means ± SEM. P-values noted determined by unpaired student t-test. (F) Beta diversity was calculated by the Bray-Curtis dissimilarity index and shown by PCoA.

After utilizing the HUMAnN 2.0 pipeline with MetaPhlAn 2.0, alpha diversity was assessed using the MicrobiomeAnalyst software package, comparing cytology brush samples to biopsy samples. Alpha diversity on the species level was found to be similar between the two groups, with higher evenness in the cytology brush group. Measures of richness included the observed richness (p=0.057, Figure 1B) and Chao1 (p=0.071, Figure 1C); measures of evenness included the Shannon (p=0.032, Figure 1D) and Simpson (p=0.055, Figure 1E) diversity measures, which were all increased in the cytology brush samples. Beta diversity was assessed as a measure of overall difference between the two samples utilizing the Bray-Curtis Index method. An overall difference was determined between the biopsy samples and the cytology brush samples (PERMANOVA, *R*^2^=0.366, F-value 4.62, p=0.028; Figure 1F). The top two axes accounted for 82.5% of diversity in the Bray-Curtis Index analysis, while the remaining axes account for 17.5%. Based upon these findings, we observed an overall trend towards increased alpha diversity in the cytology brush samples compared to biopsies (though underpowered), as well as a difference in beta diversity between the two collection methods.

### Taxonomic Profiles Minimally Differ Between Collection Methods

At the class level, the highest prevalence was found to be Actinobacteria in the biopsy sample group (56.1±21.2% versus 22.3±14.9% in the brush group, p=.02)(Figure 2A, Supplemental Table 3), while the highest prevalence in the brush group was Clostridia (47.7±10.8% versus 4.1±7.3% in the biopsy group, p=0.00007). The discordance was persistent at lower levels such as the top 10 most abundant species (Figure 2B, Supplemental Table 4), and the four statistically significantly different species as shown in Figure 2C. The top nine species were chosen as a simplistic measure of the most abundant species present and eliminating others with very low abundance. The most abundant species in the biopsy group was determined to be *Propionibacterium acnes* (53.3±18.4% versus 13.7±12.2% in the brush group, p=0.062), while the most abundant species in the brush group was *Faecalibacterium prausnitzii* (17.4±11.6% versus 2.3±3.4% in the biopsy group, p=0.12). Two additional species were found to be statistically significantly different between the two groups: *Streptococcus thermophilus* (higher in biopsy group, p=0.12) and *Ruminococcus lactaris* (higher in brush group, p=0.25). In all, there was minimal difference in terms of taxonomic profiling between the two collection methods.

**Figure 2.**
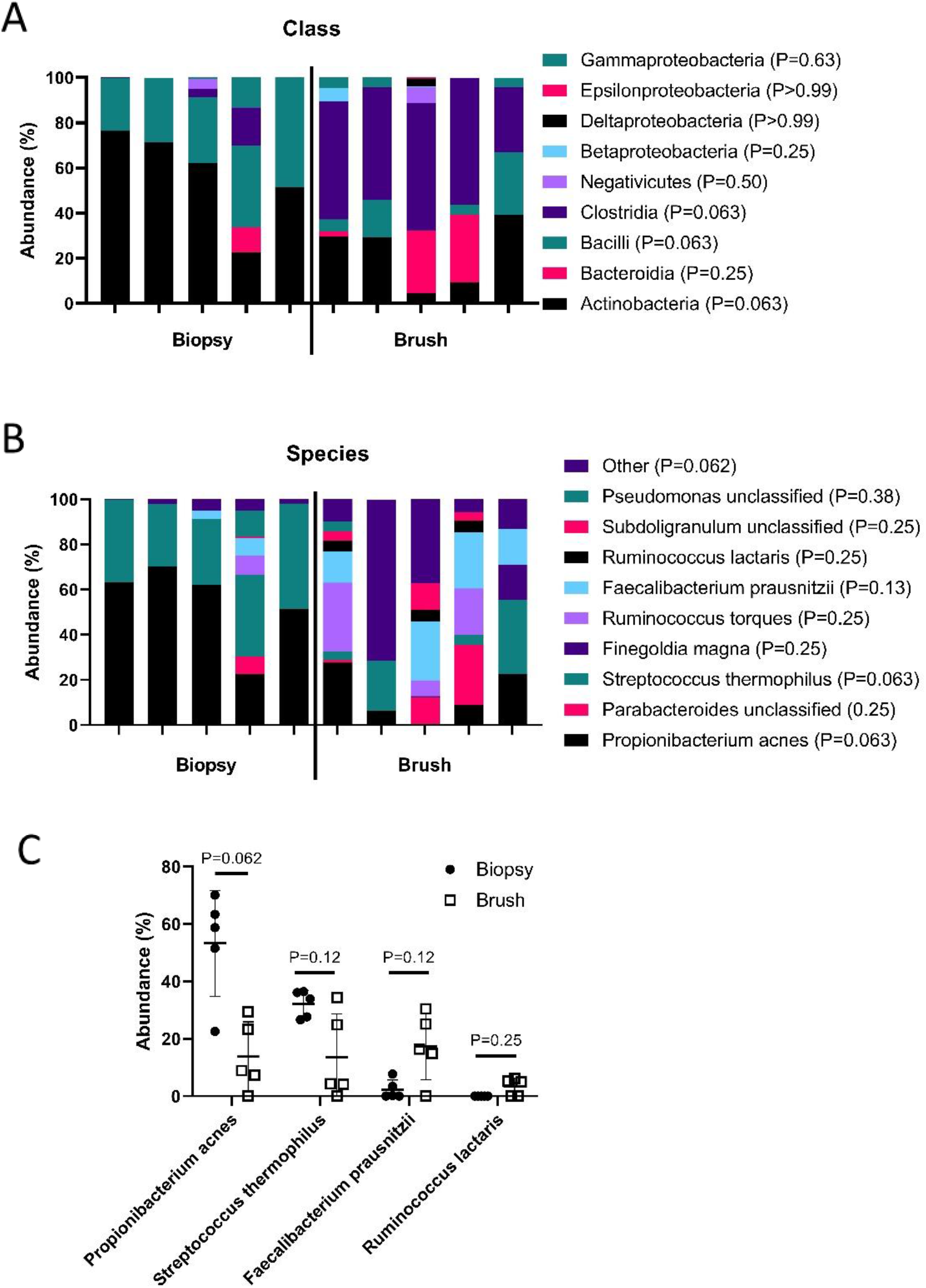
Few specific taxonomic differences are observed between collection methods. Metatranscriptomics analysis was performed on the 5 participants from each group. Biopsy and brush groups were compared with regards to (A) class and (B) species level differences, and shown as stacked bars of the mean taxa abundance within the collection method group. P-values determined by Wilcoxon signed-rank with FDR 1%. (C) The relative abundances of individual OTU are depicted as symbols for each individual, with a total of 4 different species identified as having significant differences in pair-wise comparisons. Data are shown as symbols for the abundance of the taxa for each subject, and bars as the group mean ± SEM. P-values determined by Wilcoxon signed-rank with FDR 1%.

### Metatranscriptomics Analysis Illustrates Minimal Functional Differences Between Collection Methods

Metatranscriptomics data was analyzed to assess transcribed pathway differences between the two groups. In general, overall KO metabolism pathway distribution was largely similar comparing the two groups, with no significant differences identified (Figure 3A). Within specific individual pathways, the top 10 most abundant hits were then compared (Figure 3B); two transcripts were significantly different between the two groups: K02703, photosystem II P680 reaction center D1 protein (p=0.02, higher in biopsy group), and K02961, small subunit ribosomal protein S17 (p<0.05, higher in brush group) (Figure 3C). Top 10 pathways were again analyzed as a measure of overall abundance given the significantly large numbers of pathways identified. Together, these data demonstrate minimal differences in metatranscriptomic profiling between the biopsy and cytology brush collection methods.

**Figure 3.**
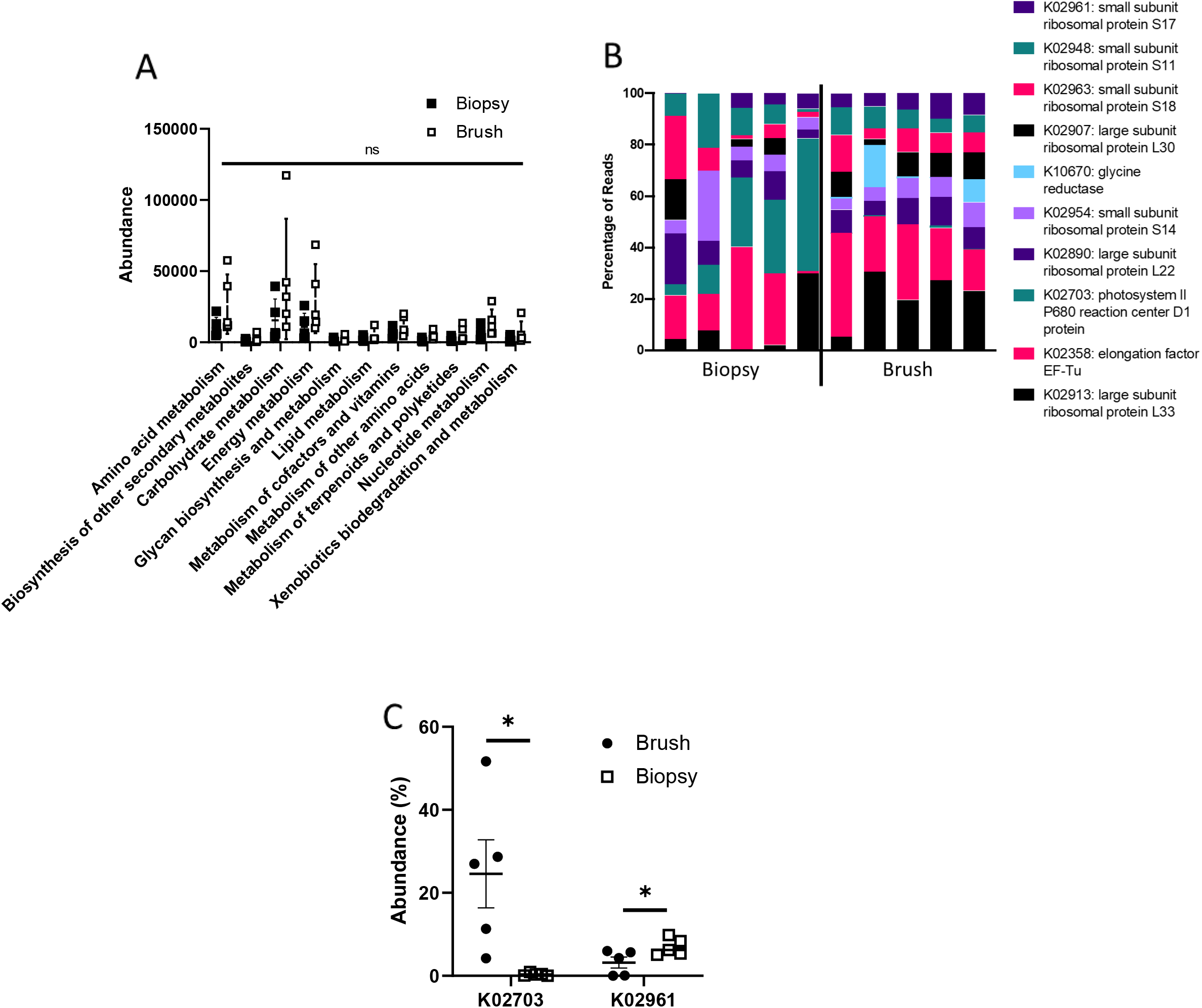
Metatranscriptomic analyses reveal minimal differences between collection methods. Metatranscriptomics data was assessed for differences in (A) overall pathways between biopsy and brush groups demonstrating no difference as determined by ANOVA with Kruskal-Wallis post-hoc test. (B) The top 10 most abundant transcriptional pathways were identified and compared between the two groups with statistical difference in K02703 and K02961 (*, p <0.05), as determined by Kruskal-Wallis post-hoc test, and demonstrated in (C) as a PCoA comparing the two different sample collection methods of biopsy versus cytology brush.

### Metabolite Profiles Diverge Between Sample Collection Methods

Given our microbiome data from cytology brush collection was similar to that from biopsies, we next sought to compare collection methods for evaluation of metabolites. In contrast, evaluation of metabolomics data revealed stark differences between the collection methods. PCoA revealed pronounced separation between the two sample methods with good intra-group correlation, although brushes demonstrated higher variability (Figure 4A). Assessment of the specific 114 metabolites profile revealed striking differences between the two collection methods (Supplemental Figure 1 and Supplemental Figure 2), which is further highlighted within the top 25 most abundant metabolites as determined by relative tissue abundance (Figure 4B). After using a fold change of 2.0 and FDR correction of 0.1, a total of 11 metabolites are identified are being significantly different based upon a volcano plot (Supplemental Figure 2, Supplemental Table 5). Unfortunately, these were largely ubiquitous metabolites in both humans and bacteria, without any bacteria-specific metabolites to compare production versus absorption. Put together, our data reveal differential findings with regards to metabolite profiles between the two sample collection methods of biopsy and cytology brush, providing an intriguing difference compared to our metatranscriptomics findings.

**Figure 4.**
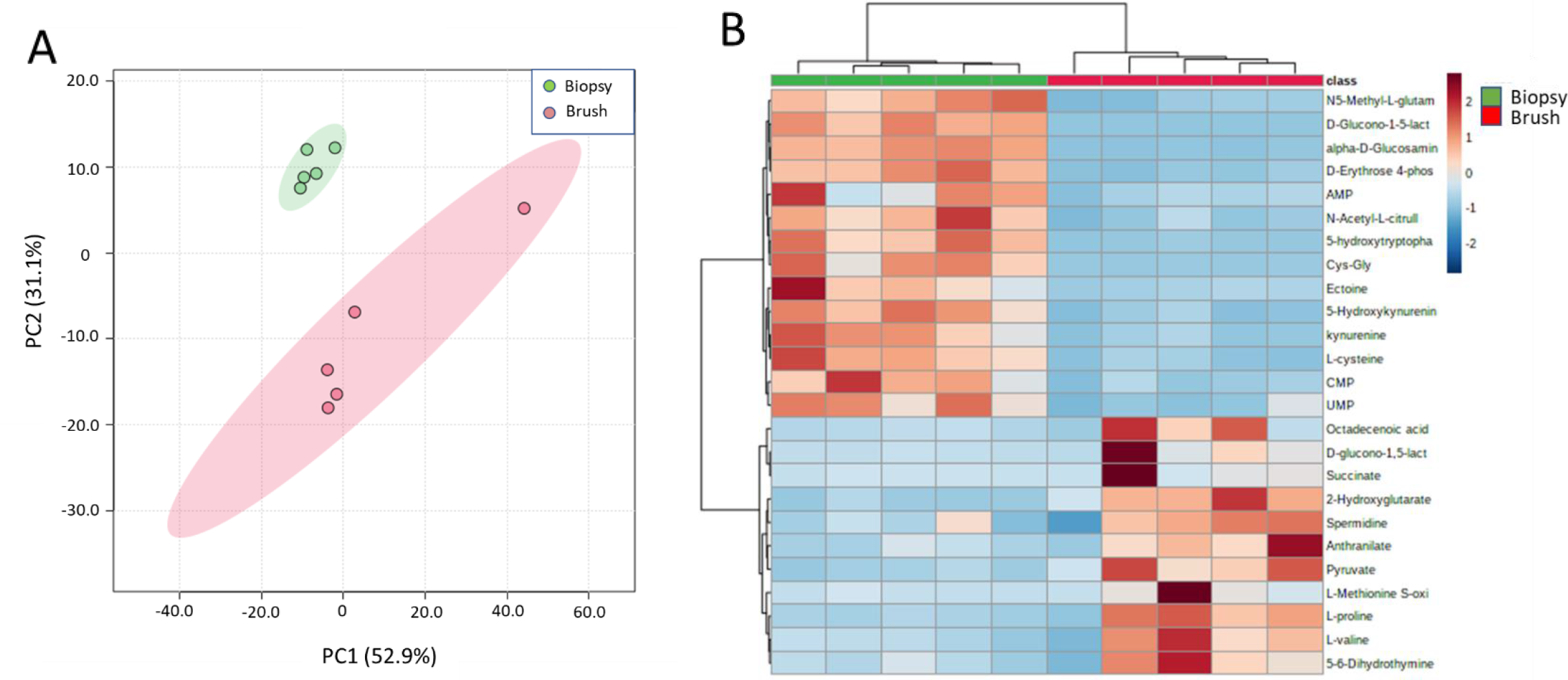
Assessment of metabolomics reveals striking differences between collection methods. Broad screening of metabolites was performed by UHPLC-MS comparing biopsies and cytology brushes. (A) PCoA of the metabolite data for all 114 identified metabolites within the two groups is shown. (B) Top 25 identified metabolites based upon t-test demonstrated as a heatmap with p<0.05 and FDR <0.05.

## Discussion

Associations between the human gut microbiome and the host immune system are characterized in numerous diseases, including autoimmune disease (23-25). The ability to study this association is predicated on appropriate methods of sample collection and ensuring samples are an accurate representation of the question at hand. In this study we start to ask the fundamental question of whether sampling techniques result in different answers based upon how collection is performed. We aim to test a new method of less invasive collection that utilizes cytology brush scrapings of the gut mucosal layer and compare it to colon pinch biopsy using a multi ‘omics approach. Our findings indicate that a similar metatranscriptomic signature can be identified, but metabolomic profile indicates very different findings, suggesting bacterially produced and human absorbed metabolites are significantly different.

One striking finding is that cytology brush scrapings of the gut mucosal layer demonstrate a significantly increased bacterial read count with transcriptomics sequencing when compared to the gold standard tissue biopsy. Our data indicates that 59.1% of the sequencing reads in the brush group correspond to bacterial nucleic acid, versus 26.5% of the biopsy group (Figure 1A). This finding suggests the low bacterial read count from gut biopsies with a much higher human nucleic acid count that must be removed for assessing the microbiome. Our cytology brush method therefore appears to better sample the mucosal microbiome without human contamination, resulting in a significantly increased bacterial nucleic acid content compared to the biopsy method. A similar result was observed in ileal pouch sampling (26). In general, alpha diversity trends towards an increase in the cytology brush group by all measures, but only reaches significance with the Shannon method. This is likely a reflection of the small sample size in this pilot study, but these findings also correlate with previous literature assessing microbial diversity across the gut mucus layer compared to the lumen(27).

With regards to beta diversity, our findings suggest greater diversity in the cytology brush group compared to the biopsy group (Figure 1F). This finding is likely a reflection of the different layers of the gut being assessed as the biopsy tends to be deeper compared to the more superficial brush which may include luminal and mucosal components. Another likely reason for this finding may be the higher bacterial content of the brush as previously described, which leads to a higher concentration of bacterially sequenced reads versus more human reads that are removed from the biopsy group.

Assessing the transcriptomics data demonstrates minor differences with regards to higher order bacterial identification as well as species identification (Figure 2). The greater microbial diversity on a species level in the brush group and less variability in the biopsy group may be due to the more superficial sampling provided by the brush as reflected by the higher alpha and beta diversity shown in Figure 1. Prior methods have attempted using similar cytology brushes during endoscopic procedures(26, 28, 29), with similar results. Other attempts to assess differences in taxonomic profiling among sampling methods have looked at fecal samples compared to colonic biopsies utilizing 16S in healthy participants, and identified differences in alpha diversity richness, beta diversity, and numerous bacterial phyla differences(30). Similarly, prior attempts to compare standard rectal swabs to biopsy have identified mixed results having similar taxonomic profiling(31, 32) versus different taxonomic profiling(33), though our study appears to be the first to attempt using a cytology brush to scrape the mucosal layer and provide transcriptomic information.

Metatranscriptomics provides functional insights regarding the microbial community, and the end products are the proteins and other biochemicals produced. In this regard, metabolomics profiling can supplement transcriptomic data. Previous studies have identified a number of different microbial-produced metabolites that influence the host immune system such as short-chain fatty acids, riboflavin metabolites, and tryptophan metabolites(34-36), which highlights the importance of methods to study this interaction. Our findings demonstrate a striking difference between sample methods in the metabolite profile as a whole (Supplemental Figure 1), as well as by PCoA and the top 25 most abundant metabolites (Figure 4). This finding is not unexpected given that different layers of the gut tissue are being analyzed - intestinal tissue with associated mucous layer in the biopsies versus the associated mucous layer itself in the brush samples. The differences in the collections highlights an important caveat in assessing the bacteria-derived metabolome: significant differences exist between metabolites that are produced versus what is absorbed, and the results of this data must be considered when critically analyzing future similar studies of gut metabolomics.

## Limitations

This study has a number of limitations that must be considered. First, as this is a pilot study for proof-of-concept, the number of participants that were recruited were low, and thus adjustment for potential confounding factors was not performed. These participants were also undergoing routine colonoscopy with bowel prep. Therefore, it is unclear how the microbiome of these participants may be different compared to other studies that rely on stool samples, as we are only assessing the mucosal layer and not all bacteria present in the gut. Repeating this study utilizing participants that have not undergone bowel prep will be of interest. The hypothesis in question relied upon metatranscriptomics analysis, which is only one potential ‘omics method for microbiome profiling, and others such as 16S sequencing or metagenomics could also be performed. Lastly, there was not a control arm in this study that included a stool sample for comparison, and this was purposely chosen due to the practical inability to perform this analysis prior to bowel prep and concern for temporal differences in sample acquisition.

## Conclusion

This pilot study aims to identify an alternate sampling method to intestinal biopsy for microbiome studies. Our results testing use of a cytology brush demonstrates the ability to sample the gut mucosal microbiome. We find that numerous different ‘omics approaches are able to be carried out through this sampling method and provides intriguing initial results to be attempted in further studies. Taxonomic profiling demonstrates minor differences while transcriptional analysis is comparable between the two methods, thus demonstrating the importance of experimental design and hypothesis generation towards sampling method for these types of studies. Metabolite assessment demonstrates a clear difference between the gut mucous layer and the tissue, and this distinction can be used for future studies that aim to understand the role of bacterially derived metabolites and how they are absorbed in the gut. In conclusion, we demonstrate early promising results of this new technique, and find new advances in the field of studying gut-microbiome interactions that provide potential for future use.

## Acknowledgements

We would like to acknowledge Francesca Cendali and Julie Reisz, Ph.D. of the University of Colorado School of Medicine Metabolomics Core for assistance with metabolomics analysis.

## Ethics Approval and Consent to Participate

This study was conducted according to the principles within the Declaration of Helsinki. All study procedures were approved by the Colorado Multiple Institutional Review Board (protocol #14-2012).

## Author Contributions

AJB and KAK conceptualized and designed the study. MEG, BPF and FIS recruited the participants. MEG, BPF and FIS acquired the samples. AJB and AB processed and extracted the samples. AJB, AS and KAK analyzed and interpreted the data. AJB, AB and KAK drafted the manuscript with input from AS. All authors contributed to the article and approved the submitted version.

## Data Availability

Raw data for the metabolomics and metagenomics sequencing will be made available upon request to the corresponding author. Sequencing data is publicly assessable in the National Library of Medicine’s Sequence Read Archive SUB 10104885.

## Figure Legends

**Supplemental Table 1.**
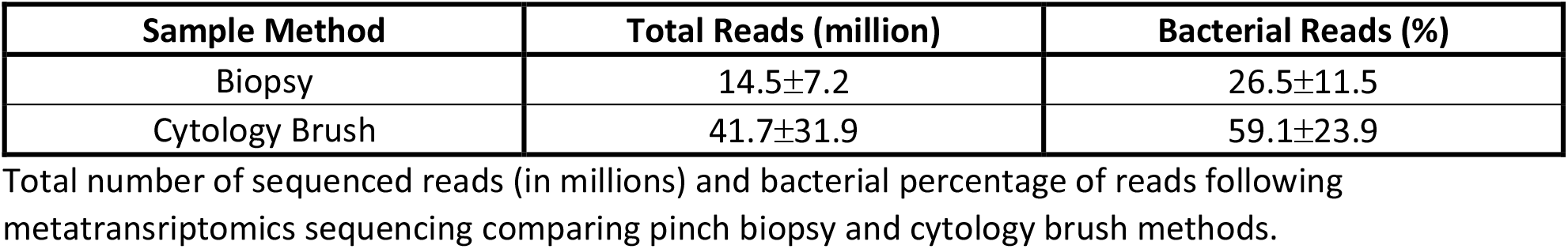

**Supplemental Table 2.**
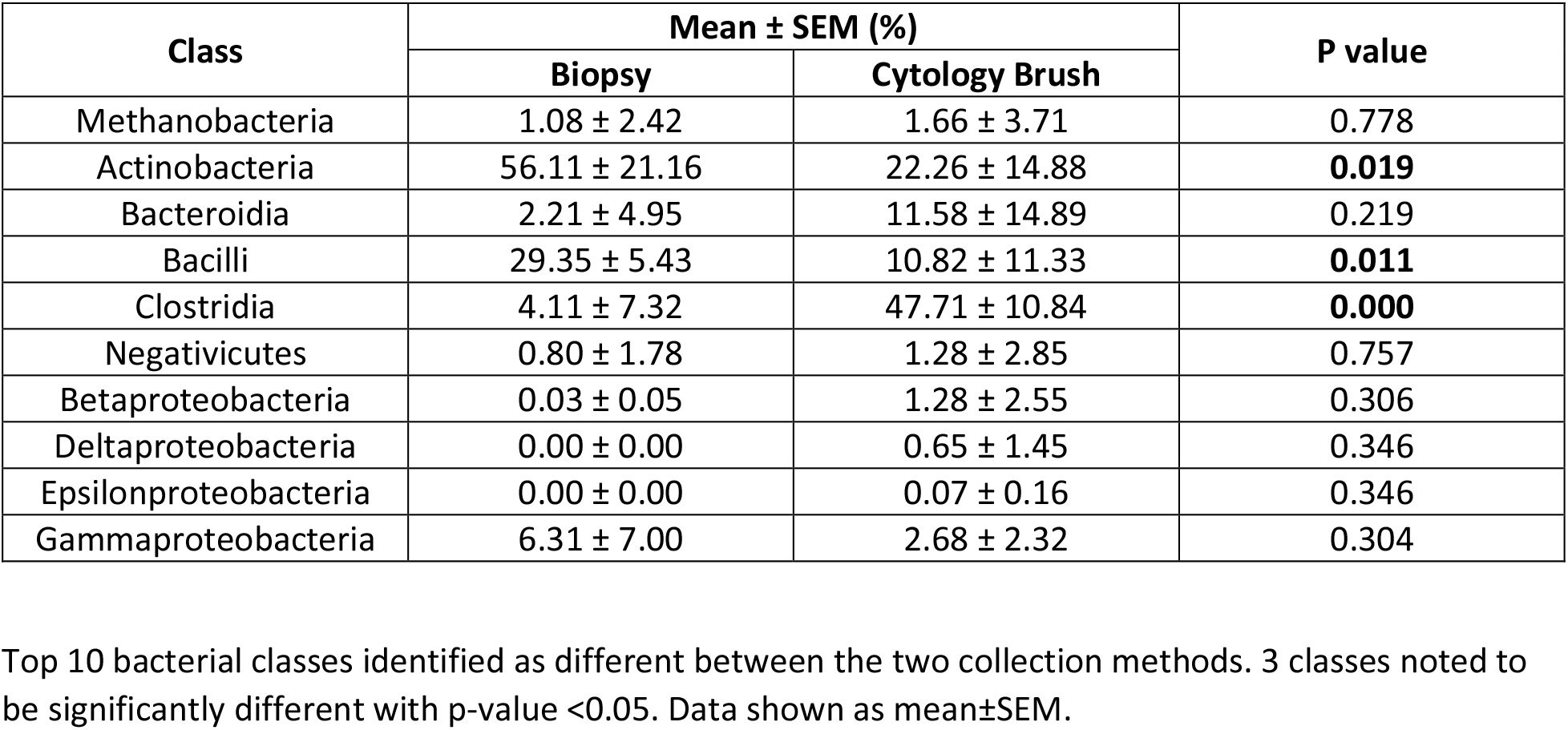

**Supplemental Table 3.**

| Species | Mean ± SEM (%) | | P-value |
| --- | --- | --- | --- |
|  | Biopsy | Cytology Brush |  |
| Faecalibacterium prausnitzii | 2.27 ± 3.36 | 17.37 ± 11.62 | **0.023** |
| Finegoldia magna | 0.00 ± 0.00 | 12.49 ± 19.77 | 0.195 |
| Methanobrevibacter unclassified | 1.08 ± 2.42 | 2.11 ± 4.72 | 0.676 |
| Parabacteroides unclassified | 1.55 ± 3.46 | 8.52 ± 11.94 | 0.246 |
| Propionibacterium acnes | 53.2 ± 18.44 | 13.76 ± 12.17 | **0.004** |
| Pseudomonas unclassified | 5.13 ± 6.44 | 0.93 ± 2.07 | 0.202 |
| Ruminococcus lactaris | 0.01 ± 0.02 | 3.27 ± 3.02 | **0.042** |
| Ruminococcus torques | 1.72 ± 3.84 | 12.22 ± 14.01 | 0.145 |
| Streptococcus thermophilus | 32.14 ± 4.69 | 13.54 ± 15.18 | **0.031** |
| Subdoligranulum unclassified | 0.13 ± 0.29 | 4.35 ± 5.54 | 0.127 |
| Others | 2.68 ± 2.09 | 11.44 ± 7.71 | **0.039** |

Top 10 bacterial species identified as different between the two collection methods. 4 species noted to be significantly different with p-value <0.05. Data shown as mean ± SEM.

**Supplemental Figure 1.**
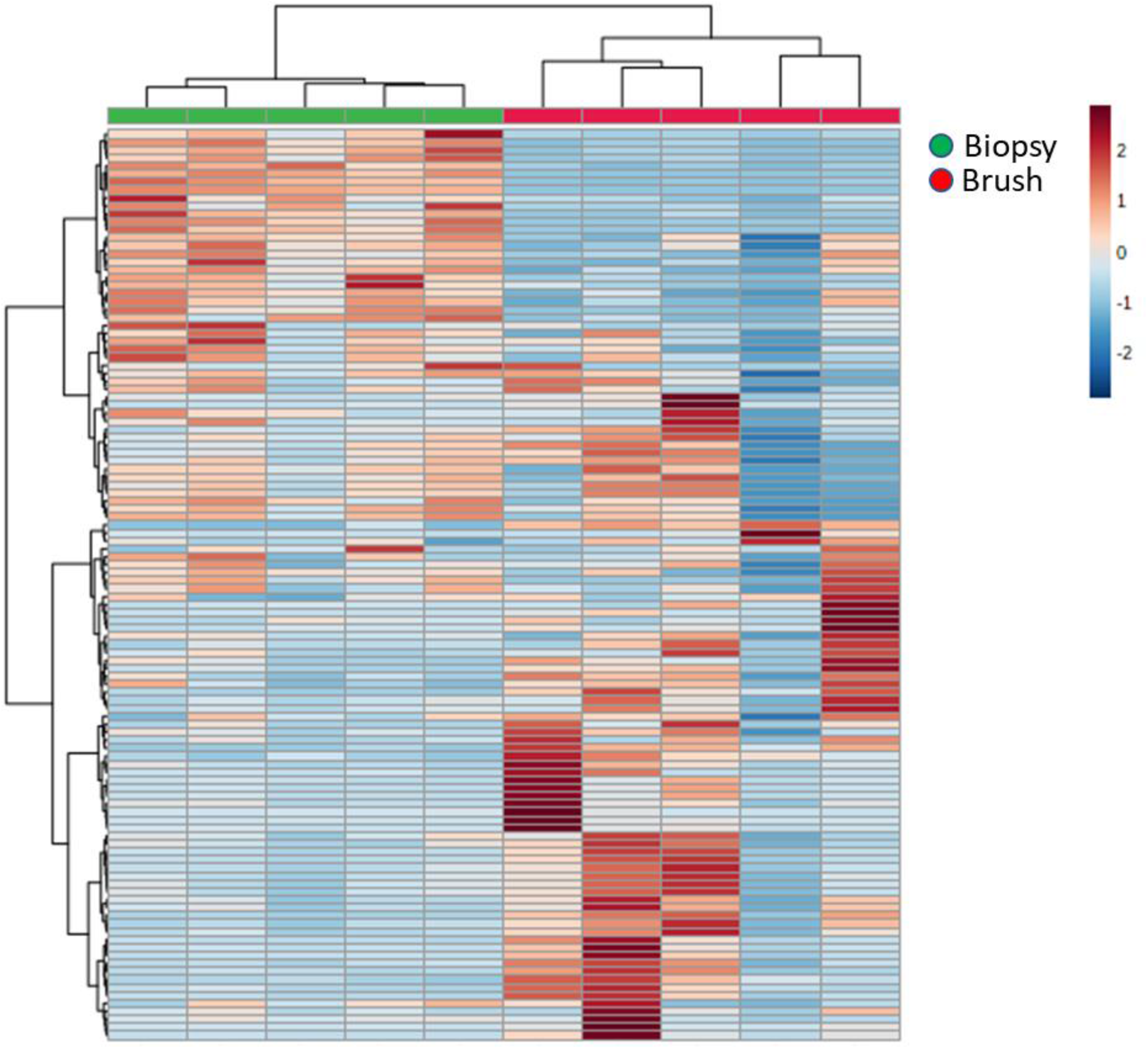
Heat map demonstrating all 114 metabolites comparing the pinch biopsy versus cytology brush collection methods.

**Supplemental Figure 2.**
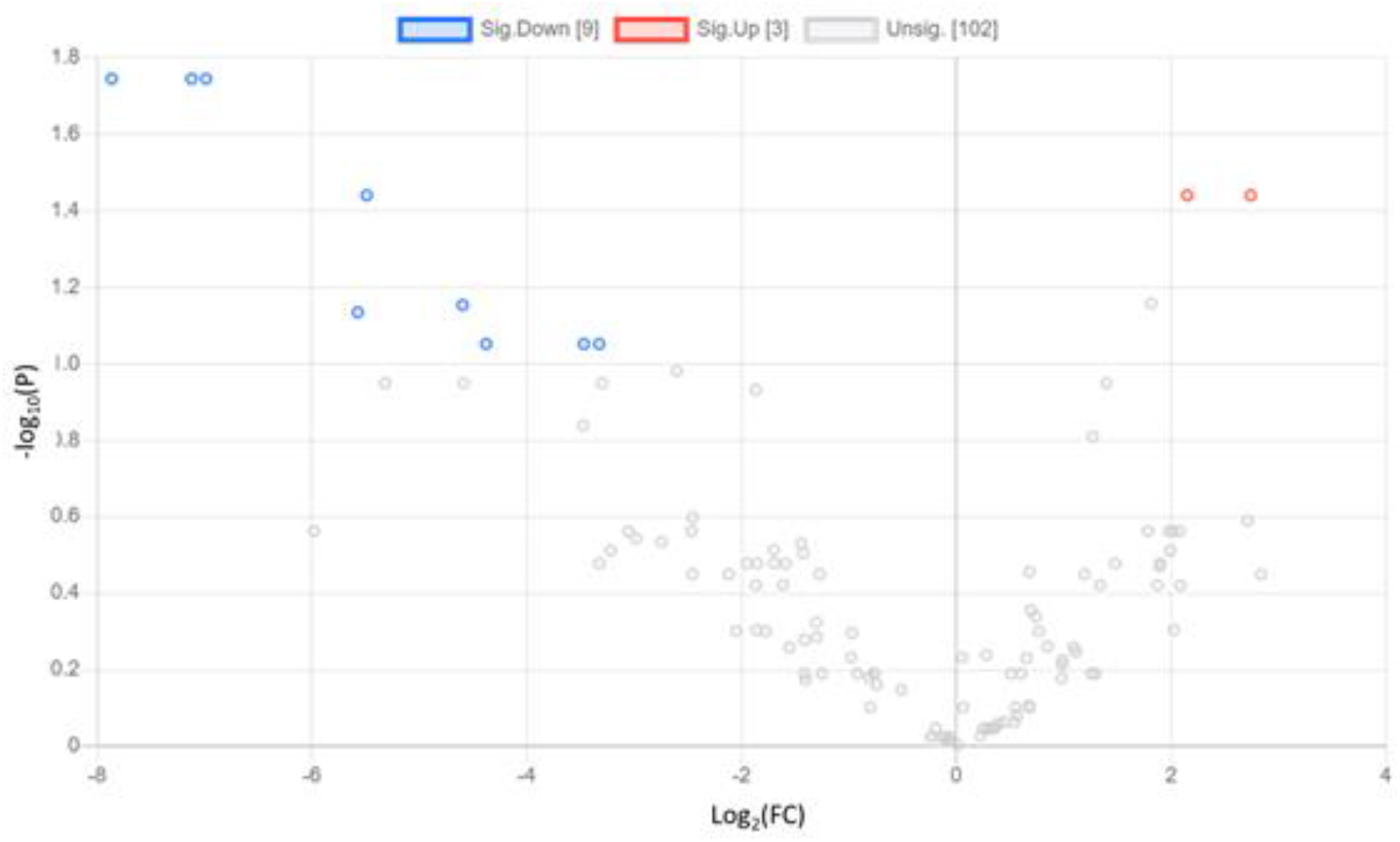
Volcano plot demonstrating 114 metabolites identified as significantly up (red), down (blue), or unchanged (gray) comparing the pinch biopsy versus cytology brush collection method. Significance determined by fold change >2.0 and Wilcoxon signed-rank test with FDR adjusted p-value <0.1.

**Supplemental Table 4.**

| Metabolite | Log<sub>2</sub>(FC) | P-value (FDR adjusted) |
| --- | --- | --- |
| D-Glucono-1-5-lactone 6-phosphate | -7.86 | 0.018 |
| Cys-Gly | -7.12 | 0.018 |
| alpha-D-Glucosamine 1-phosphate | -6.98 | 0.018 |
| 5-hydroxytryptophan | -5.45 | 0.036 |
| 2-Hydroxyglutarate/Citramalate | 2.74 | 0.036 |
| Itaconate | 2.15 | 0.036 |
| L-cysteine | -4.59 | 0.070 |
| D-Erythrose 4-phosphate | -5.57 | 0.073 |
| N5-Methyl-L-glutamine | -4.38 | 0.089 |
| N-Acetyl-L-citrulline | -3.47 | 0.089 |
| CMP | -3.32 | 0.089 |

Metabolites identified as significant (fold change, FC >2.0 and paired-t-test with FDR adjusted p-value <0.1) for comparison between pinch biopsy and cytology brush collection method.

## References

1. Lozupone CA, Stombaugh JI, Gordon JI, Jansson JK, Knight R. Diversity, stability and resilience of the human gut microbiota. Nature. 2012;489(7415):220–30.

2. Weiss GA, Hennet T. Mechanisms and consequences of intestinal dysbiosis. Cell Mol Life Sci. 2017;74(16):2959–77.

3. Levy M, Blacher E, Elinav E. Microbiome, metabolites and host immunity. Curr Opin Microbiol. 2017;35:8–15.

4. Levy M, Kolodziejczyk AA, Thaiss CA, Elinav E. Dysbiosis and the immune system. Nat Rev Immunol. 2017;17(4):219–32.

5. Zhang X, Li L, Butcher J, Stintzi A, Figeys D. Advancing functional and translational microbiome research using meta-omics approaches. Microbiome. 2019;7(1):154.

6. Donaldson GP, Lee SM, Mazmanian SK. Gut biogeography of the bacterial microbiota. Nat Rev Microbiol. 2016;14(1):20–32.

7. Sheth RU, Li M, Jiang W, Sims PA, Leong KW, Wang HH. Spatial metagenomic characterization of microbial biogeography in the gut. Nat Biotechnol. 2019;37(8):877–83.

8. Tang Q, Jin G, Wang G, Liu T, Liu X, Wang B, et al. Current Sampling Methods for Gut Microbiota: A Call for More Precise Devices. Front Cell Infect Microbiol. 2020;10:151.

9. Durbán A, Abellán JJ, Jiménez-Hernández N, Ponce M, Ponce J, Sala T, et al. Assessing gut microbial diversity from feces and rectal mucosa. Microb Ecol. 2011;61(1):123–33.

10. Rangel I, Sundin J, Fuentes S, Repsilber D, de Vos WM, Brummer RJ. The relationship between faecal-associated and mucosal-associated microbiota in irritable bowel syndrome patients and healthy subjects. Aliment Pharmacol Ther. 2015;42(10):1211–21.

11. Fung TC, Artis D, Sonnenberg GF. Anatomical localization of commensal bacteria in immune cell homeostasis and disease. Immunol Rev. 2014;260(1):35–49.

12. Hansson GC. Role of mucus layers in gut infection and inflammation. Curr Opin Microbiol. 2012;15(1):57–62.

13. Zoetendal EG, von Wright A, Vilpponen-Salmela T, Ben-Amor K, Akkermans AD, de Vos WM. Mucosa-associated bacteria in the human gastrointestinal tract are uniformly distributed along the colon and differ from the community recovered from feces. Appl Environ Microbiol. 2002;68(7):3401–7.

14. Bolger AM, Lohse M, Usadel B. Trimmomatic: a flexible trimmer for Illumina sequence data. Bioinformatics. 2014;30(15):2114–20.

15. Langmead B, Salzberg SL. Fast gapped-read alignment with Bowtie 2. Nat Methods. 2012;9(4):357–9.

16. Franzosa EA, McIver LJ, Rahnavard G, Thompson LR, Schirmer M, Weingart G, et al. Species-level functional profiling of metagenomes and metatranscriptomes. Nat Methods. 2018;15(11):962–8.

17. Segata N, Waldron L, Ballarini A, Narasimhan V, Jousson O, Huttenhower C. Metagenomic microbial community profiling using unique clade-specific marker genes. Nat Methods. 2012;9(8):811–4.

18. Chong J, Liu P, Zhou G, Xia J. Using MicrobiomeAnalyst for comprehensive statistical, functional, and meta-analysis of microbiome data. Nat Protoc. 2020;15(3):799–821.

19. Dhariwal A, Chong J, Habib S, King IL, Agellon LB, Xia J. MicrobiomeAnalyst: a web-based tool for comprehensive statistical, visual and meta-analysis of microbiome data. Nucleic Acids Res. 2017;45(W1):W180-W8.

20. Gehrke S, Rice S, Stefanoni D, Wilkerson RB, Nemkov T, Reisz JA, et al. Red Blood Cell Metabolic Responses to Torpor and Arousal in the Hibernator Arctic Ground Squirrel. J Proteome Res. 2019;18(4):1827–41.

21. Nemkov T, Reisz JA, Gehrke S, Hansen KC, D’Alessandro A. High-Throughput Metabolomics: Isocratic and Gradient Mass Spectrometry-Based Methods. Methods Mol Biol. 2019;1978:13–26.

22. Xia J, Wishart DS. Web-based inference of biological patterns, functions and pathways from metabolomic data using MetaboAnalyst. Nat Protoc. 2011;6(6):743–60.

23. Asquith MJ, Stauffer P, Davin S, Mitchell C, Lin P, Rosenbaum JT. Perturbed Mucosal Immunity and Dysbiosis Accompany Clinical Disease in a Rat Model of Spondyloarthritis. Arthritis Rheumatol. 2016;68(9):2151–62.

24. Knights D, Lassen KG, Xavier RJ. Advances in inflammatory bowel disease pathogenesis: linking host genetics and the microbiome. Gut. 2013;62(10):1505–10.

25. Zhang X, Zhang D, Jia H, Feng Q, Wang D, Liang D, et al. The oral and gut microbiomes are perturbed in rheumatoid arthritis and partly normalized after treatment. Nat Med. 2015;21(8):895–905.

26. Huse SM, Young VB, Morrison HG, Antonopoulos DA, Kwon J, Dalal S, et al. Comparison of brush and biopsy sampling methods of the ileal pouch for assessment of mucosa-associated microbiota of human subjects. Microbiome. 2014;2(1):5.

27. Lavelle A, Lennon G, O’Sullivan O, Docherty N, Balfe A, Maguire A, et al. Spatial variation of the colonic microbiota in patients with ulcerative colitis and control volunteers. Gut. 2015;64(10):1553–61.

28. Shanahan ER, Zhong L, Talley NJ, Morrison M, Holtmann G. Characterisation of the gastrointestinal mucosa-associated microbiota: a novel technique to prevent cross-contamination during endoscopic procedures. Aliment Pharmacol Ther. 2016;43(11):1186–96.

29. Nishino K, Nishida A, Inoue R, Kawada Y, Ohno M, Sakai S, et al. Analysis of endoscopic brush samples identified mucosa-associated dysbiosis in inflammatory bowel disease. J Gastroenterol. 2018;53(1):95–106.

30. Carstens A, Roos A, Andreasson A, Magnuson A, Agréus L, Halfvarson J, et al. Differential clustering of fecal and mucosa-associated microbiota in ‘healthy’ individuals. J Dig Dis. 2018;19(12):745–52.

31. Jones RB, Zhu X, Moan E, Murff HJ, Ness RM, Seidner DL, et al. Inter-niche and inter-individual variation in gut microbial community assessment using stool, rectal swab, and mucosal samples. Sci Rep. 2018;8(1):4139.

32. Budding AE, Grasman ME, Eck A, Bogaards JA, Vandenbroucke-Grauls CM, van Bodegraven AA, et al. Rectal swabs for analysis of the intestinal microbiota. PLoS One. 2014;9(7):e101344.

33. Araújo-Pérez F, McCoy AN, Okechukwu C, Carroll IM, Smith KM, Jeremiah K, et al. Differences in microbial signatures between rectal mucosal biopsies and rectal swabs. Gut Microbes. 2012;3(6):530–5.

34. Kelly CJ, Zheng L, Campbell EL, Saeedi B, Scholz CC, Bayless AJ, et al. Crosstalk between Microbiota-Derived Short-Chain Fatty Acids and Intestinal Epithelial HIF Augments Tissue Barrier Function. Cell Host Microbe. 2015;17(5):662–71.

35. Magalhaes I, Solders M, Kaipe H. MAIT Cells in Health and Disease. Methods Mol Biol. 2020;2098:3–21.

36. Hubbard TD, Murray IA, Bisson WH, Lahoti TS, Gowda K, Amin SG, et al. Adaptation of the human aryl hydrocarbon receptor to sense microbiota-derived indoles. Sci Rep. 2015;5:12689.

